# The human brain views selfish behaviour towards genetic vs. non-genetic sibling differently

**DOI:** 10.1101/112383

**Authors:** Mareike Bacha-Trams, Enrico Glerean, Juha Lahnakoski, Elisa Ryyppö, Mikko Sams, Iiro P. Jääskeläinen

**Author notes:** Correspondence to: Mareike Bacha-Trams and Iiro P. Jaaskelainen, Department of Neuroscience and Biomedical Engineering, School of Science, Aalto University, P.O. Box 12200, FI-00076 AALTO, Finland.

## Abstract

Previous behavioural studies have shown that humans act more altruistically towards kin. Whether and how such kinship preference translates into differential neurocognitive evaluation of social interactions has remained an open question. Here, we investigated how the human brain is engaged when viewing a moral dilemma between genetic vs. non-genetic sisters. During functional magnetic resonance imaging, a movie depicting refusal of organ donation between two sisters was shown, with participants guided to believe the sisters were related either genetically or by adoption. The participants selfreported that genetic relationship was not relevant to them, yet their brain activity told a different story. When the participants believed that the sisters were genetically related, inter-subject similarity of brain activity was significantly stronger in areas supporting response-conflict resolution, emotion regulation, and self-referential social cognition. Our results show that mere knowledge of a genetic relationship between interacting persons can robustly modulate social cognition of the perceiver.

## Introduction

Imagine watching a movie wherein a healthy teenage girl refuses to donate a kidney to her fatally ill sister, thus letting her die. The healthy sister even sues her parents, challenging their plea to help her sister by arguing for her right to physical integrity. Would you feel that the healthy sister was doing the right thing? Then imagine knowing that the sisters are not related genetically but that the sister who is asked to donate was adopted at young age. Would this knowledge affect your perception of the organ refusal? It has been demonstrated in previous behavioural studies that kinship matters when making altruistic decisions^1,2,3,4,5^, of which donating one’s kidney can be considered as an extreme case.

Evaluating and predicting social interactions of others is an integral part of social cognition, one of the most important of human cognitive functions. For instance, the evolution of social cognition may best explain why humans have a more developed neocortex than other species^6^. So far, social cognition has been predominantly studied with stimuli depicting interactions between strangers, however, a great deal if not majority of significant interactions evaluated in daily life are between one’s family members, friends, and acquaintances. In many cultures, including the Finnish culture where there is great respect for private space of others, social interactions are typically limited to situations where a person either meets others that he/she knows from beforehand, or to specific scenarios where interactions with strangers are needed to achieve a specific goal, e.g., asking a salesperson for help when buying a product. Behavioral studies have shown that the evaluation of morality of actions differs for genetically related and unrelated others^4,5^. Further, previous neuroimaging studies have shown that prior knowledge about protagonists can lead to differential processing when observing their actions: brain activity differed when subjects thought that the person performing actions were either trustworthy or not trustworthy, or whether the harm described was intentional or accidental^7,8,9^. Here, we set forth to investigate whether kin are perceived differently based on prior knowledge of presence vs. absence of a genetic relationship.

We studied this question with functional magnetic resonance imaging (fMRI): 30 healthy female participants watched an edited 24-min version of the movie “My sister’s keeper” (dir. Nick Cassavetes, 2009, Curmudgeon Films) depicting a girl refusing to donate a kidney to her sister suffering from cancer. Due to improvements in fMRI acquisition methods and data analysis algorithms^10^ it has become possible to study specific aspects of social cognition between participants using ecologically valid fMRI paradigms. When viewing a feature film during brain scanning, not only basic sensory cortices but also “higher-order” prefrontal cortical areas become synchronized across participants^11,12^. Therefore, we calculated inter-subject correlations of voxel time courses to reveal areas participating in processing the movie content.

Each participant watched the same movie four times under different conditions: In two scanning sessions on two separate days four to six weeks apart, the participants were informed prior to viewing, that the sisters are either genetically related or that the sister who is expected to donate her organ was adopted at a young age. Within each scanning session they also watched the movie after being asked to adopt the perspective of either the to-be-organ-donor or the perspective of the sister who is supposed to receive the organ (for the sake of focus, results related to taking different perspectives are not reported here). The case of sisters related genetically vs. by adoption is highly suitable for testing the perception of a moral dilemma among kin given that, unlike in case of e.g. in-laws, there is no potential for shared genetic interest in future generations for adopted siblings^13^. Should knowledge of genetic relationship matter, we hypothesize that brain regions known to be involved in processing of mentalising^14,15^, conflict resolution^16,17^, emotion regulation^18,19^, and moral dilemmas ^20,21^ would be activated differently under the two viewing conditions.

To evaluate specifically which brain areas are associated with the perception and processing of moral dilemmas during movie watching, each participant underwent a second experiment performing moral decision task during fMRI scanning. In the task the participants had to decide to save individuals from a dangerous area and had different choices including their own sister, best friend and strangers. Again, if the genetic relationship had an effect on the viewers, we hypothesize that that the participants show kinship preference by saving their sister over others and that similar brain areas are engaged in the decision task and when watching the movie believing that the sisters are genetic.

Further, to complement the neuroimaging data, additional physiological and psychological data were collected. Heart rate, breathing rate, and eye-movements were recorded during fMRI. After the fMRI, the subjects were asked whether genetic vs. non-genetic relationship status mattered to them in the moral dilemma that they observed. They were also re-shown the clips and asked to rate continuously, on separate runs, the emotional valance and arousal that they had experienced when they viewed the movie during fMRI.

The subjects self-reported that genetic vs. non-genetic status of the sisters was not relevant to them. Eye-movements, heart rate, breathing rate, and self-ratings of emotional valance and arousal were also highly similar between these two conditions. In contrast, there were significant differences in ISC in several brain regions between the genetic vs. non-genetic viewing conditions. Areas of significantly higher synchrony in the case of viewing genetic rather than non-genetic sisters included the dorsolateral prefrontal cortex (DLPFC), the ventromedial prefrontal cortex (VMPFC), the inferior frontal gyrus (IFG) the cuneus and precuneus, the anterior and posterior cingulate (ACC and PCC), the superior parietal lobule (SPL), the superior temporal sulcus and gyrus (STS/STG) and the Insula, which have been in previous literature associated with supporting response-conflict resolution, emotion regulation, and self-referential social cognition. These areas overlapped with those activated in the separate moral dilemma decision task. Thus, the results suggest that that mere knowledge of a genetic relationship between interacting persons can robustly modulate social cognition of the perceiver.

## Results

### Inter-subject correlation (ISC) across all conditions

In a first step, the overall ISC^22^ of hemodynamic activity of all participants (N = 30) was calculated during first viewing of the movie. The resulting ISC map (Figure 1) thus includes all the conditions equally. Significant ISC was observed extensively in occipital lobes, posterior parietal areas, and temporal cortices. In the frontal cortex, areas in the lateral inferior frontal gyrus, lateral medial frontal gyrus, DLPFC, dorsomedial prefrontal cortex (DMPFC) and VMPFC showed ISC between all participants.

**Figure 1.**
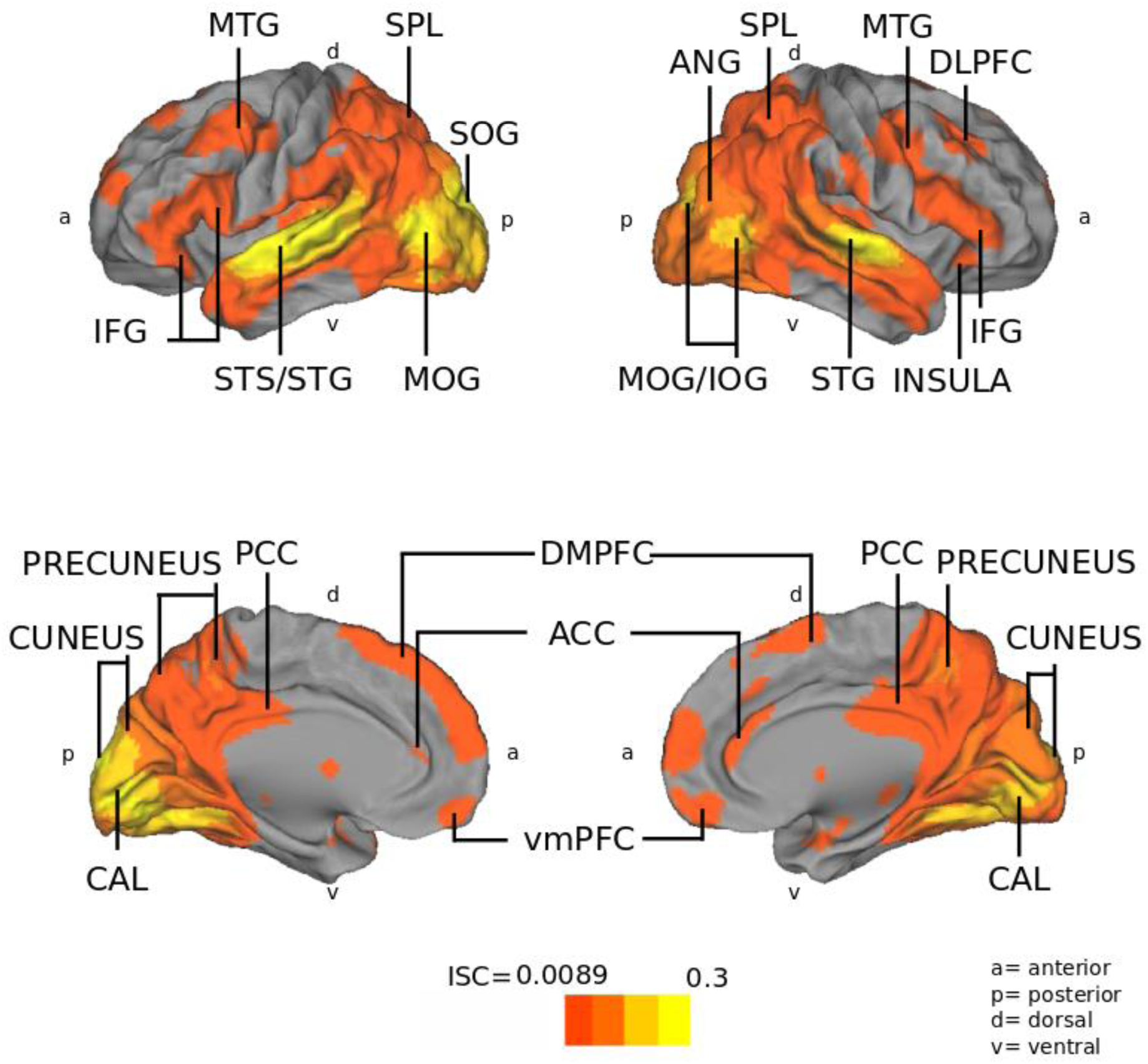
Inter-subject correlation (ISC) of all 30 participants during the first viewing of the movie. On top row are shown lateral and on bottom row medial surfaces of left and right cerebral hemispheres. Red-yellow colours indicate areas of significant ISC during movie watching (FDR q< 0.05). Abbreviations: ACC = anterior cingulate cortex, CAL = calcarine gyrus, DLPFC = dorsolateral prefrontal cortex IFG = inferior frontal gyrus, IOG = inferior occipital gyrus, MFC = medial frontal cortex, MOG = medial occipital gyrus, MTG = medial temporal gyrus, PCC = posterior cingulate cortex. SPL = superior parietal lobe, STG/STS = superior temporal gyrus and sulcus, VMPFC = ventromedial prefrontal cortex.

### Differences in ISC between conditions

In a second step the ISC of all participants were contrasted between the genetic vs. non-genetic relationship viewing conditions. In a behavioural questionnaire, 90% of the participants selfreported that it did not matter to them whether the sisters were related genetically or by adoption. In contrast, when the participants watched the movie believing to see genetically related sisters, the ISC were significantly stronger in the STS/STG, VMPFC, DLPFC, ACC and PCC, IFG, insula, cuneus, precuneus, and SPL (Figure 2, Table S1). When the participants thought the sisters were non-genetic, higher ISC was observed mainly in the occipital cortex. Importantly, this robustly differential neural processing of the movie was triggered solely by different knowledge of the genetic status of the sisters, as the movie stimulus *per se* was identical in both viewing conditions.

**Figure 2.**
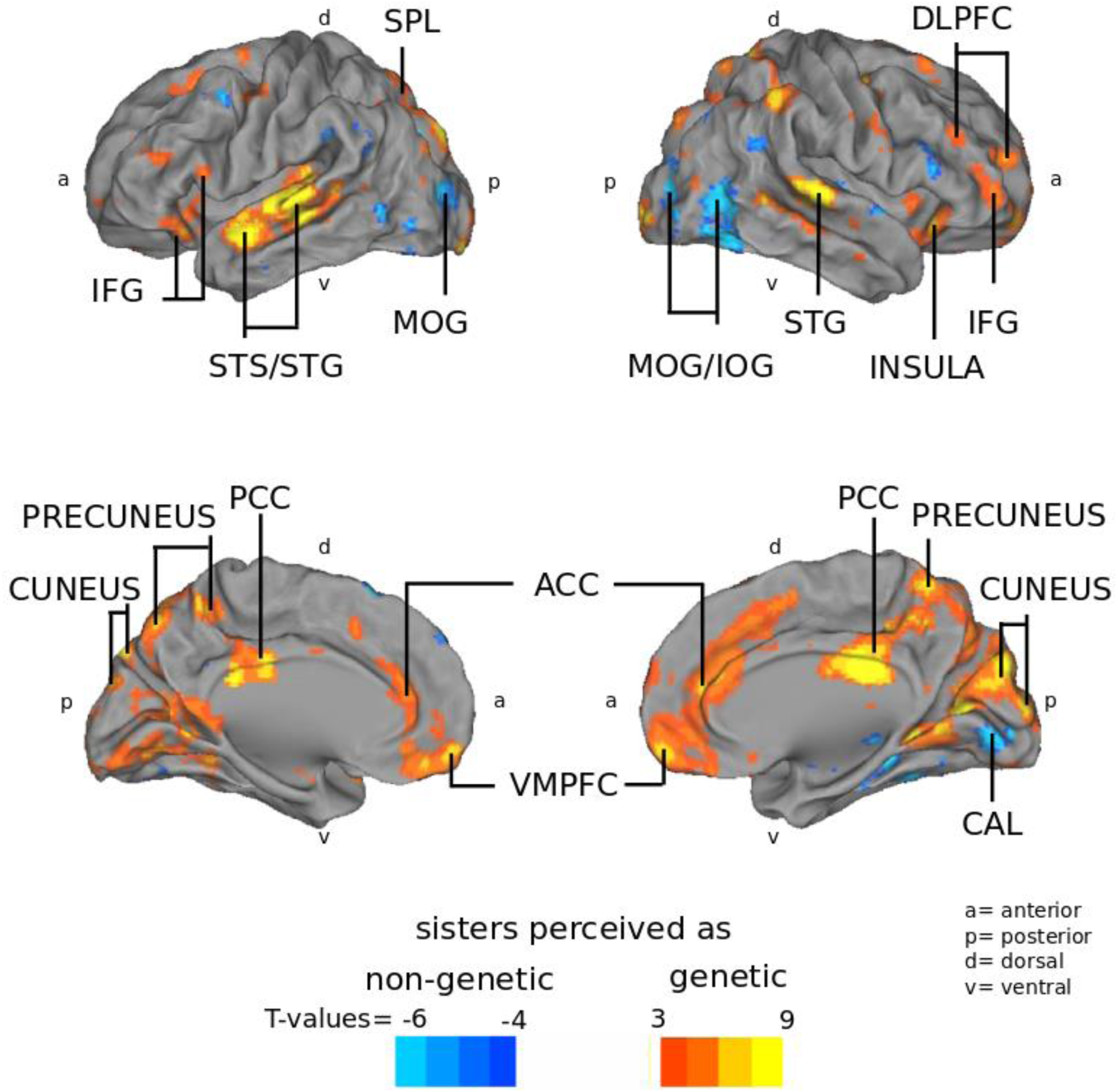
Significant differences in brain activity when all 30 participants watched the movie thinking that the sisters were genetically vs. non-genetically related (FDR q< 0.05). Red-yellow colours indicate areas of significantly higher ISC when the participants watched the movie as depicting genetically related, as compared to non-genetically related, sisters. Blue colour indicates areas showing significantly higher ISC in the reverse contrast. Warm-coloured areas include areas known to support moral judgement (DLPFC, precuneus, STS/STG), conflict resolution and emotion regulation (ACC, Insula), self-referential social cognition (VMPFC), as well as mentalizing (SPL, STS/STG, precuneus and PCC). Blue-coloured areas have been associated with perception of visual information (MOG/IOG,CAL).

One possible explanation for these findings is that there might be a general implicit bias against adopted sisters that modulates brain functions during movie viewing. To test for this possibility, we used an implicit association test (IAT^23^) in an independent behavioural control experiment (N=30). Reaction times during assignment of positive and negative connoted words to the categories of sister and adopted sister showed that there is no such implicit bias: Out of 30 participants, nine favoured a genetic sister, 13 a non-genetic sister, and eight had no preference (one sampled t-test t = −0,9564 p= 0,3468 non-significant). The TOST procedure^24^ indicated that the ratings of emotional closeness were significantly similar (observed effect size (d = −0,17) was significantly within the equivalent bounds of d = −0,68 and d = 0,68, t(29) = 2,77, p = 0,005).

Furthermore, eye-movements, heart rate and breathing rate, recorded simultaneously with fMRI, did not show any significant differences between the conditions. Self-ratings of emotional valence and arousal obtained after the scans were not significantly different between these two conditions (Valence: r = 0.0075, p = 0.3458, Arousal: r = −0.0189, p = 0.6081) (Figure S1).

### Comparison with the moral dilemma decision control experiment

To further examine which neurocognitive processes could be involved, we studied whether the brain areas showing higher ISC in the genetic condition overlap with those engaged during a modified moral dilemma task^25^ (Figure S2, Table S2). When the same participants who underwent the movie viewing experiment had to decide between saving their sister, best friend, vs. stranger(s), in various combinations, from crisis regions, 93% of the participants showed kinship preference, by choosing their sister rather than their friend (chi-squared test Xi^2^ = 43.4 p< 4 × 10^−11^). Further, as can be seen in Figure 3 the VMPFC, DLPFC, ACC, PCC, precuneus, IFG, MTG, SPL and anterior insula were consistently involved in both making choices when one’s kin is preferred over others and when observing the refusal of organ donation to a genetic vs. non-genetic sister. Importantly, ratings of emotional closeness were not significantly different between the participants’ sisters and their best friends with an average of 9.28 (sisters) and 8.80 (friends) on a 1-10 scale (Wilcoxon signed rank test = 0.12, t-test, t = 1.64 p = 0.11). The TOST procedure^24^ indicated that the ratings of emotional closeness were significantly similar (observed effect size (d = 0,33) was significantly within the equivalent bounds of d = −0,68 and d = 0,68, or in raw scores: −1,07 and 1,07, t(29) = −1,92, p = 0,032.

**Figure 3.**
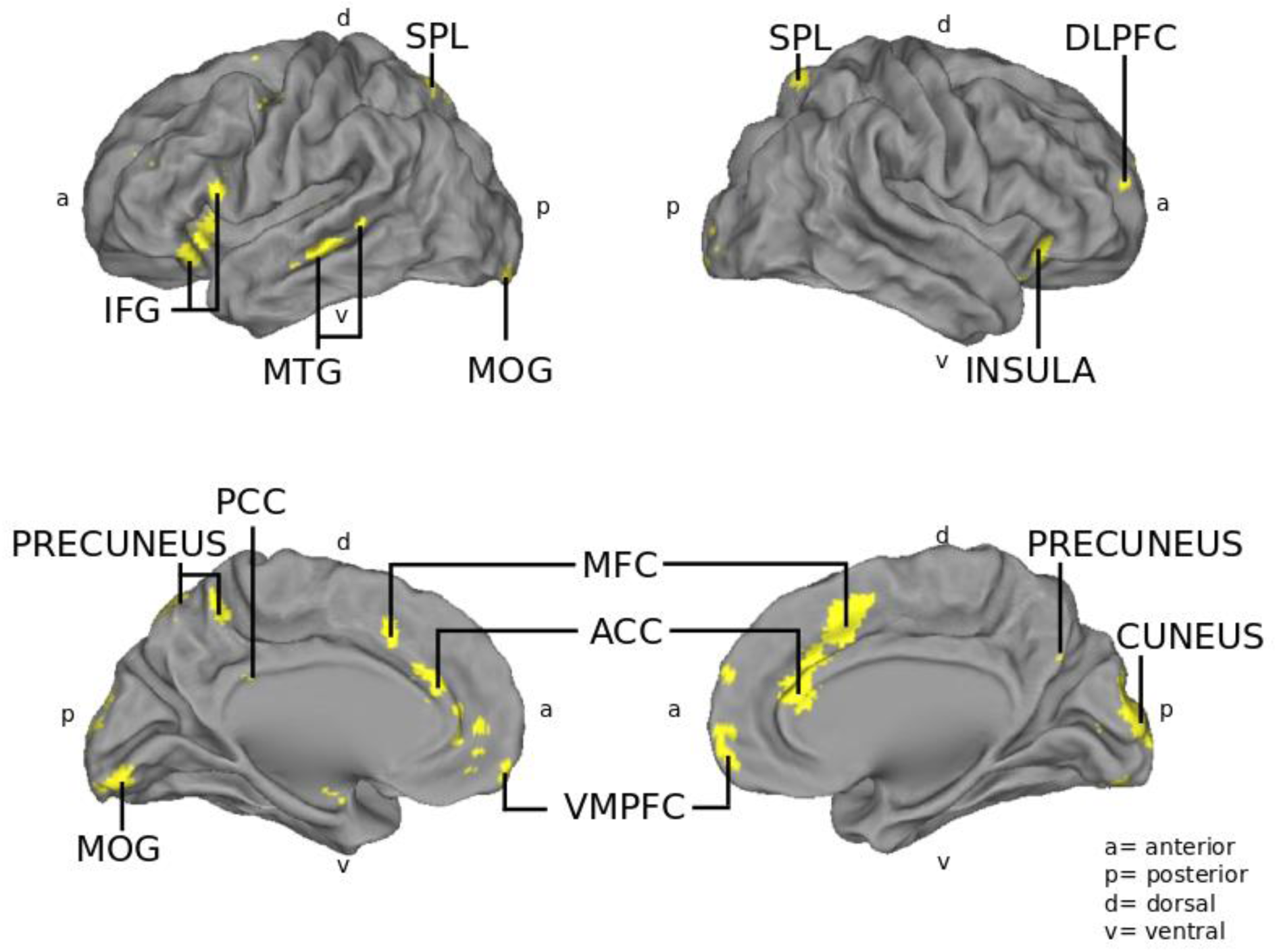
Yellow colour indicates brain areas where there was both higher ISC during watching the movie while assuming the sisters to be genetically related and significant hemodynamic activity during the moral kinship decision task, N = 30. Brain area labels as in Figure1.

**Figure 4.**
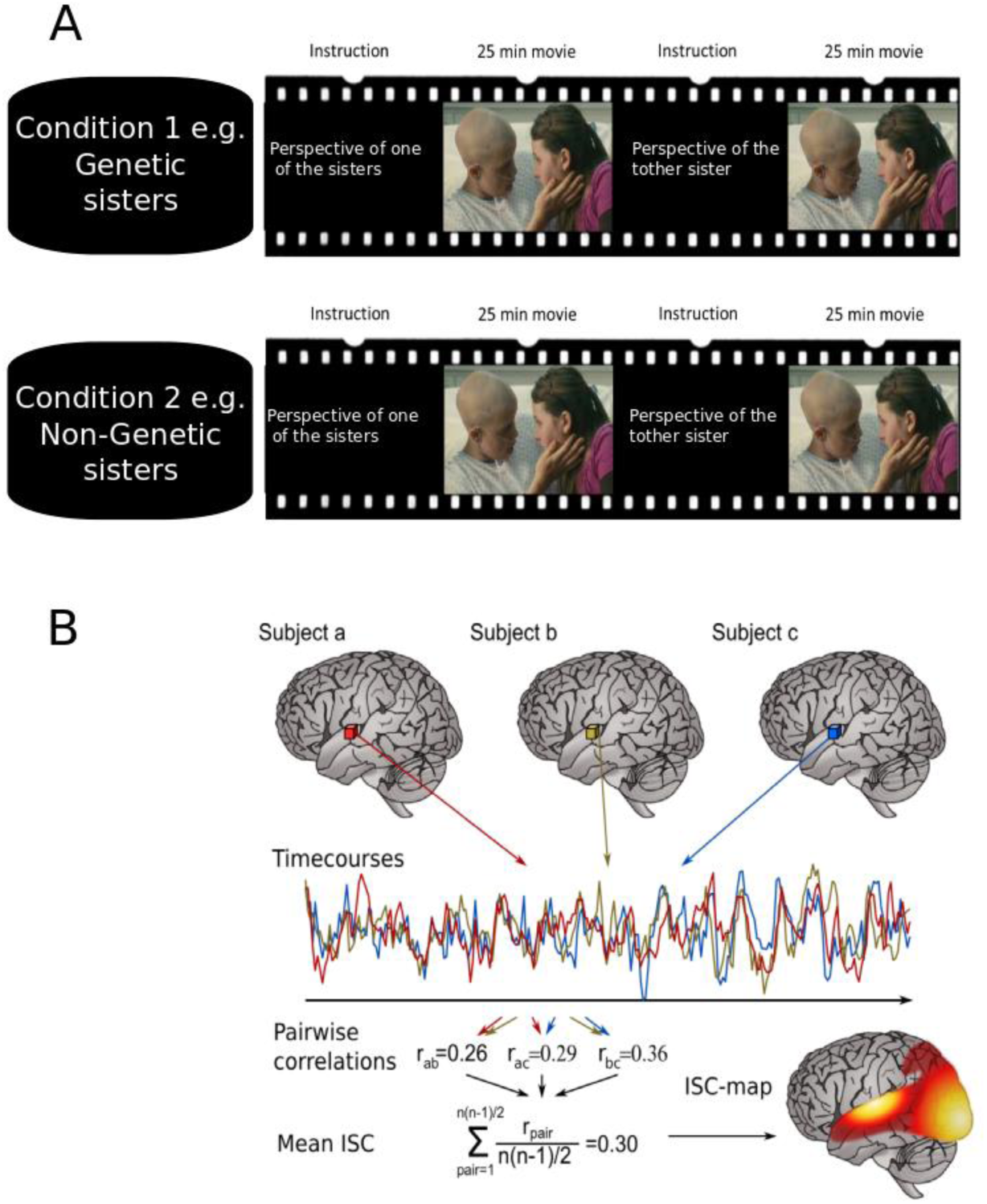
Experimental procedure and ISC analysis in the movie watching task. A: Every participant watched the movie four times, in a 2 × 2 design assuming that the movie characters are either genetic sisters or not genetically related and taking the perspective of the to-be-donor sister Anna or the to-be-recipient sister Kate. The order of all the conditions were counter-balanced. B: Time series from each voxel from all the fMRI recordings are compared across participants in pairwise correlations to obtain the mean inter-subject-correlation (ISC).

## Discussion

In the present study, we investigated whether refusing altruism from a sister is perceived differently when the viewers think that the sisters are genetically related vs. when they think that one of the sisters has been adopted at young age. Indeed, there were multiple brain regions showing significantly higher ISC when the subjects thought that they are seeing a young girl refusing to donate her organ to save her genetic, as opposed, to non-genetic, sister (Figure 2). These areas included VMPFC, DLPFC, ACC, PCC, insula, precuneus, and SPL. In previous studies, these areas have been shown to be associated with moral and emotional conflict regulation^26^, decision making^27^, mentalising^20^, and perspective taking^28^, thus suggesting more uniform engagement of such neurocognitive functions when observing the dilemma of organ donation between genetic sisters. When the participants thought the sisters were non-genetic, higher ISC was observed in brain areas in the occipital cortex, which has been previously associated mainly with visual perception^29,30^. One possible explanation could potentially be that in case of a non-genetic relationship between the sisters, the processing of complex social conflict of the moral dilemma is less demanding, and leaves therefore room for the participants to focus on the visual aspects of the movie. However, eye-movements were not significantly different between the two viewing conditions, suggesting that differences in attention to movie events did not explain the observed robust differences in ISC.

In a separate experiment, we ruled out the possibility that these results could be explained by a general implicit bias to favour genetic over adopted sisters. The IAT^23^, applied to test for this possibility, showed no bias in any direction. Further, heart rate and breathing rate with no significant differences, as well as similar self-reported emotional valance and arousal between the conditions (Figure S1), in turn suggests that there were no robust differences in experienced emotions between the conditions that could have explained the observed differences in ISC. Thus, the differences in ISC between the conditions appear to have arisen due to the knowledge about the sisters’ relationship influencing cognitive evaluation of the moral dilemma depicted in the movie.

To specifically test for the possibility that the differences in ISC between the conditions reflected differences in moral evaluation, we compared the areas showing differences in ISC with areas activated when the subjects engaged in a separate moral-decision making control task. In this control task, the participants had to make choices of saving people from crisis regions including their sister, best friend, and strangers, in various combinations. Significantly longer reaction times suggested increased difficulty when having to choose between one’s sister and best friend. Areas in the VMPFC, DLPFC, ACC, PCC, precuneus, IFG, MTG, SPL and anterior insula were consistently involved in both making choices where one’s kin is preferred over others and when observing the refusal of organ donation to a genetic vs. non-genetic sister (Figure 3). This overlap in engaged brain regions suggests that processing of moral dilemmas took place during movie watching when the sisters were perceived as genetically related.

Keeping in mind the caveats associated with reverse inference^31^ (although see^32^), of these areas the DLPFC has been reported to play a role in overcoming a primary moral judgment in favour of greater welfare^14,27^ and in cortical emotional processing^33^,the MTG has been implicated in attributing mental states as well as ingroup-outgroup distinctions^21,34,35^, the ACC has been reported to be engaged in resolving conflicts^16,17^, and the SPL, precuneus, and PCC have been implicated in mentalising and perspective taking^15,36,37^. Further, the VMPFC has been associated with viewing moral conflicts, making moral decisions, attributing mental states to self and others, adopting another person’s perspective, and evaluating their beliefs^19,28,37,38^.

In summary, we observed robust differences in brain activity when participants viewed an identical movie depicting refusal to donate an organ to a genetic vs. non-genetic sister. These differences in brain activity were observed despite the participants self-reporting that the relational status of the sisters did not make any difference to them. Areas of increased synchrony in the case of genetic sisters overlapped with those activated in a separate moral dilemma decision task. Taken together, our results suggest that the precuneus, MTG, insula, SPL, and the VMPFC, along with the associated cognitive processes (i.e., moral and emotional conflict regulation, decision making, mentalising, and perspective taking) are synchronized across participants more robustly when they are viewing refusal of altruism from genetic, as opposed to non-genetic, kin. Overall, these findings are fundamentally important for understanding social cognition, a pivotal ability that makes us humans and enables, among other things, the existence of societies. Our findings point out that the perceived relationship of interacting persons robustly modulates how the brains of spectators process those interactions. This is highly significant given that vast majority of research has been on social cognition of strangers, whereas a wealth, if not majority, of significant interactions take place between one’s family members, friends, and acquaintances.

## Material and Methods

### Participants

We studied 33 healthy female participants^39^ (19-39 years, mean age of 26 years, one lefthanded, laterality index of right handed 84.5%). None of the participants reported any history of neurological or psychiatric disorders. When asked, all participants reported either normal vision or corrected to normal vision by contact lenses. Three participants were excluded due to discomfort in the scanner, so that the final analysis included 30 participants. 27 of them were native Finnish speakers and three were native Russian speakers. All participants were sufficiently proficient in English to follow the dialogue in the movie without subtitles. The study was carried out with the permission from the research ethics committee of the Aalto University and in accordance with the guidelines of the declaration of Helsinki^40^. Written informed consent was obtained from each participant prior to participation.

### Stimuli and Procedure

The study consisted of two experiments. In the first experiment, the feature film “My sister’s keeper” (dir. Nick Cassavetes, 2009, Curmudgeon Films), edited to 23 minutes and 44 secs with the main story line retained, was shown to the participants during fMRI. This shortened version of the movie focuses on the moral dilemma of the protagonist Anna to donate one kidney to her sister Kate, who is fatally ill from cancer. In the course of the movie Anna refuses to donate and Kate dies. The reason for Anna refusing to donate the kidney was not revealed to the participants until after the experiment. The movie was shown to the participants in the scanner four times in two separate scanning sessions on two different days. For each viewing of the movie the instructions were varied regarding the information about the sister’s relationship and the perspective to take in this viewing. Each participant thus watched the movie assuming that the sisters were genetic sisters or that the younger sister Anna had been adopted as a newborn. In addition each participant was asked to take either the perspective of the potential donor (Anna) or the perspective of the potential recipient (Kate) on separate viewings (and both under the condition of a genetic or non-genetic relation background). The order of the different viewing conditions was counterbalanced between the participants.

In the second, control, experiment, each participant carried out a moral-dilemma decision task during fMRI in order to localize brain regions that are related to moral decision making. For this purpose, a modified version of the classical trolley dilemma ^25,41,42^ was shown to the participants. Each participant had to choose between rescuing different individuals, including unknown individuals, their sister, and best female friend. A presentation showing text and pictures told a story about civil unrest in a distant fictive country. This country had two parts: a very dangerous and a little less dangerous. Different people are in both parts of the country. It was further told that as the participant is a very rich person, owning an airplane, she could go there and rescue some of the people. Due to the circumstances in the country she has to decide, however, which group of people to rescue. The two choices were always a group of five individuals on one side and a single person on the other. In seven runs the identity of the involved individual(s) was varied using the real names of the participant’s sister and best female friend. The 7 runs were: 1. All persons are unknown, 2. Sister is with four others in the dangerous part of the country, the single person is unknown, 3. Five persons are in the dangerous part of the country, the single person is the sister, 4. Five persons are in the dangerous part of the country, the single person is the friend, 5. Sister is with four others in the dangerous part of the country, the single person is the friend, 6. Friend is with four others in the dangerous part of the country, the single person is the sister, 7. Sister is with four other in the less dangerous part of the country, the single person is unknown. Responses in the moral dilemma decision task were recorded with button press with a LUMItouch keypad (Photon Control Inc.8363, Canada).

### fMRI acquisition

Before each scan the participants were informed about the scanning procedures and asked to avoid bodily movements during the scans. All stimuli were presented to the participant with the Presentation software (Neurobehavioral Systems Inc., Albany, CA, USA), synchronizing the onset of the stimuli with the beginning of the functional scans. The movie was back-projected on a semitransparent screen using a data projector (PT-DZ8700/DZ110X Series, Panasonic, Osaka, Japan). The participants viewed the screen at 33-35 cm viewing distance *via* a mirror located above their eyes. The audio track of the movie was played to the participants with a Sensimetrics S14 audio system (Sensimetrics Corporation Malden, USA). The intensity of the auditory stimulation was individually adjusted to be loud enough to be heard over the scanner noise. The brain-imaging data were acquired with a 3T Siemens MAGNETOM Skyra (Siemens Healthcare, Erlangen, Germany), at the Advanced Magnetic Imaging center, Aalto University, using a standard 20-channel receiving head-neck coil. Anatomical images were acquired using a T1-weighted MPRAGE pulse sequence (TR 2530 ms, TE 3.3 ms, TI 1100 ms, flip angle 7°, 256 x 256 matrix, 176 sagittal slices, 1-mm^3^ resolution). Whole-brain functional data were acquired with T2*-weighted EPI sequence sensitive to the BOLD contrast (TR 2000 ms, TE 30 ms, flip angle 90, 64 × 64 matrix, 35 axial slices, slice thickness 4 mm, 3×3 mm in plane resolution). A total of 712 whole-brain EPI volumes were thus acquired for each movie viewing of a session. The number of whole-brain EPI volumes for the moral dilemma decision task varied individually due to the involved decision made by each participant (median 267 whole-brain EPI volumes).

Heart pulse and respiration were monitored with the Biopac system (Biopac Systems Inc., Isla Vista, California, USA) during fMRI. Instantaneous values of heart rate and breathing rate were estimated with Drifter software package^43^ (http://becs.aalto.fi/en/research/bayes/drifter/).

### Recording of eye-movements

Eye movements were recorded during fMRI scanning from all participants with an EyeLink 1000 eye tracker (SR Research, Mississauga, Ontario, Canada; sampling rate 1000 Hz, spatial accuracy better than 0.5°, with a 0.01° resolution in the pupil-tracking mode). Due to technical problems, 4 participants had to be excluded from the final data analysis (with the rejection criteria of blinks maximum 10% of the duration of the scan and majority of blinks and saccades less than 1 second of duration). Prior to the experiment the eye tracker was calibrated once with a nine-point calibration.

Saccade detection was performed using a velocity threshold of 30°/s and an acceleration threshold of 4000°/s2. Because the experiment was relatively long and no intermediate drift correction was performed, we retrospectively corrected the mean effect of the drift. We first calculated the mean of all fixation locations over the entire experiment for each participant, and then rigidly shifted the fixation distributions so that the mean fixation location coincided with the grand mean fixation location over all participants.

### Behavioral Measurements and Self-reports

#### Valence and Arousal measurements

The participants self-reported emotions they had experienced during movie viewing. This was carried out after the fMRI experiment by viewing the movie again (Full procedures have been described in an earlier publication^44^). Two aspects of emotional experience were rated: emotional valence (positive-negative scale) and arousal which were acquired on separate runs. While watching the movie in the middle of the screen, the participants could move a small cursor on the right side of the screen up and down on a scale using the computer mouse to report their current state of valence or arousal using a web tool https://version.aalto.fi/gitlab/eglerean/dynamicannotations^44^. The self-ratings were collected at 5 Hz sampling rate.

#### Behavioral questionnaires

The participants were also asked after the first fMRI session five short freeform questions about their perception of the movie, specifically about how easy it was to take one or the other perspective, and whether they would have donated their kidney if in place of the movie protagonist. After the second fMRI session all participants were debriefed by showing them the ending of the original movie, where it is revealed that the sick sister had wished for the healthy sister to refuse donating her kidney. Afterwards they were asked if seeing the real ending changed their opinion on the roles of the two movie protagonists.

As an additional self-report measure, the participants’ disposition for catching emotions from others was assessed with two emotional empathy questionnaires: Hatfield’s Emotional Contagion Scale^45^ and the BIS/BAS scale^46^. Every participant also filled in a questionnaire quantifying their social network ^47^, including their emotional closeness to their sister and best friend. The names of the sister and best friend were obtained from this questionnaire for the moral dilemma task.

#### Perspective taking

In the movie-viewing experiment, in addition to having the participants to watch the movie in the conditions of sisters related by birth or by adoption, we had altogether four runs, so that on two of the runs the participants were asked to view the movie from the perspective of the sister who was expected to donate the organ, and on two of the runs from the perspective of the to-be-recipient sister. Thus, there was one run wherein the participants viewed the movie from the perspective of the to-be-donor thinking that the sisters were genetic, one run wherein the participants viewed the movie from the perspective of the to-be-donor thinking that the sisters were non-genetic, one run wherein the participants viewed the movie from the perspective of the to-be-recipient thinking that the sisters were genetic, and one run wherein the participants viewed the movie from the perspective of the to-be-recipient thinking that the sisters were nongenetic.

### fMRI preprocessing

Standard fMRI preprocessing steps were applied using the FSL software (www.fmrib.ox.ac.uk) and custom MATLAB code (available at https://version.aalto.fi/gitlab/BML/bramila/). Briefly, EPI images were corrected for head motion using MCFLIRT and coregistered to the Montreal Neurological Institute 152 2mm template in a two-step registration procedure using FLIRT: from EPI to participant’s anatomical image after brain extraction (9 degrees of freedom) and from anatomical to standard template (12 degrees of freedom). Further, spatial smoothing was applied with a Gaussian kernel of 6 mm full width at half maximum. High pass temporal filter at a cut-off frequency of 0.01 Hz was used to remove scanner drift. To further control for motion and physiological artefacts, BOLD time series were cleaned using 24 motion-related regressors, signal from deep white matter, ventricles and cerebral spinal fluid locations (see^48^ for details, cerebral spinal fluid mask from SPM8 file csf.nii, white matter and ventricles masks from Harvard Oxford atlas included with FSL).

### Inter-subject correlation (ISC) analysis of brain activity during movie watching

To investigate how similar the brain activity was across participants in the different experimental conditions, we performed inter-subject correlation (ISC) using the isc-toolbox (https://www.nitrc.org/projects/isc-toolbox/)^49^. For each voxel the toolbox computes a similarity matrix between participant pairs and within same participant in all conditions, with the conditions being: (i) shared assumption that the movie’s sisters are genetically related, (ii) shared assumption that the younger sister was adopted, (iii) shared perspective of the to-be-donor Anna, and (iv) shared perspective of the to-be-recepient Kate. The total size of the similarity matrix is then 120×120 (4 conditions × 30 participants). Each value of the correlation matrix is a result of the correlation between the BOLD time series of the pair of participants considered for the selected voxel. We computed differences between experimental conditions by first transforming the correlation values into z-scores with the Fisher Z transform and then computing t-values and corresponding p-values using a permutation based approach^50^. Correction for the multiple comparison was performed with Benjamini-Hochberg false discovery rate (BH-FDR) correction at a q < 0.05, corresponding to a t-value threshold of 2.133. For visualization purposes, all results were also cluster corrected by removing any significant cluster smaller than 4 × 4 × 4 voxels. Summary tables were generated with an increased t-value threshold of 3. Unthresholded statistical parametric maps can be found in neurovault: http://neurovault.org/collections/WGSQZWPH/

### General linear model analysis of the fMRI data acquired during the control task

A moral dilemma decision task was performed by all participants to localize regions involved in moral processing. The moral dilemma decision task was analyzed with a general linear model approach using the SPM12 software (www.fil.ion.ucl.ac.uk/spm). To distinguish between moments of decision in the moral dilemma and the simple perception of the presentation, we created a temporal model of the occurrence of decision moments during the experiment. The decision regressor included time points from the revelation of the identity of involved individuals to the moment of decision indicated by button press. The activity during these time points was compared to the activity in all other time points of the task, including telling the background story of the moral dilemma in the presentation. Regressors were convolved with canonical hemodynamic response function to account for hemodynamic lag. From the preprocessed input data (see above) low-frequency signal drifts were removed by high-pass filtering (cutoff 128 s). First, individual contrast images were generated for the main effects of the regressors, then first level analyses were subjected to second-level analyses in MATLAB using one-sample *t*-test to test which brain areas showed significant activations in decision vs. no decision moments in a one-sample *t*-test over participants. Statistical threshold was set at *p* < 0.05 (cluster-corrected using the threshold free cluster enhancement approach implemented by FSL randomize with 5000 permutations).

### Eye-movement analysis

Participant-wise gaze fixation distributions were compared across the genetic vs. non-genetic conditions in the movie viewing task. Individual heat maps were generated by modelling each fixation as a Gaussian function using a Gaussian kernel with a standard deviation of 1degree of visual angle and a radius of 3 standard deviations. The heat maps were generated in time windows of 2 seconds corresponding to the TR used in the fMRI measurements. Spatial similarities between each pair of heat maps across the eye-tracking sessions were calculated using Pearson’s product-moment correlation coefficient (inter-subject correlation of eye gaze, eyeISC^51^). In the end a similarity matrix was obtained with correlations between each pair for each of the 712 time windows.

First, the mean eISC scores over all 712 time windows were examined. These mean scores were acquired by extracting the mean of Fisher’s Z-transformed correlation scores and then transforming these mean values back to the correlation scale before the statistical analysis. The statistical significance of the group differences was analysed by contrasting pairs in which both participants assumed a genetic relationship with pairs in which both participants assumed the younger sister to be adopted. Non-parametric permutation tests with a total of 100000 permutations were used to avoid making assumptions about the data distribution. In this procedure the data were mixed randomly to change groupings and differences in the resulting new randomised groups were used to form an estimated distribution of the data. A comparison of how many of the permuted random partitions into groups build a more extreme group mean difference that the one observed with the original grouping yielded the final p-values.

### Analysis of behavioral measurements

#### Valence and arousal measurements

To test whether dynamic valence and arousal were different between the genetic and nongenetic condition, we first computed inter-subject similarity matrices using valence and arousal rating time-series. These were compared against a similarity matrix for the experimental conditions of the viewing preceding the valence/arousal rating, i.e. the model tests for the case where individuals are more similar within the same condition (genetic or non-genetic), but dissimilar between conditions. Tests were performed using Mantel test with 5000 permutations. We also performed a test to see if subjects who were rating arousal and valence for the genetic condition had a stronger group similarity than subjects who rated arousal and valence for the non-genetic condition. Tests were performed using permutation based t-tests, As dynamic ratings can also be different in specific time points, we also performed a permutation-based t-test on valence and arousal values at each time point corrected for multiple comparisons across time.Heart rate and breathing rate analysis

Differences between experimental conditions were computed in the same way as in the ISC analysis: Correlation values were first transformed into z-scores with the Fisher Z’s transform and then a permutation based approach was used to compute t-values and corresponding p-values^51^. Correction for the multiple comparisons was performed with Benjamini-Hochberg false discovery rate (BH-FDR) correction at a q < 0.05, corresponding to a t-value threshold of 2.133.

#### Behavioral measurements with a new group of participants

Subsequent to the fMRI experiments a new group of 30 participants (all female, and having a sister, 18-33 years, mean age 25.5 years, right handed) were recruited for two further behavioral measurements. The participants first performed an implicit association test (IAT). The IAT measures attitudes and beliefs that might not be consciously self-recognized by the participant or attitudes that the participants are unwilling to report. By asking the participants to sort, as quickly as possible, positively and negatively connoted words into categories, the IAT can measure the reaction times of the association process between the categories and the evaluations (e.g., good, bad). It has been shown in previous studies that making a response is easier and thus faster if the category is matching the implicit evaluation the participant bears in mind^22^. In this study the two categories were “genetic sister” (sisko) and “adopted sister” (adoptiosisko). The two categories were paired in different randomized runs with positive or negative words, thus the experiment comprised separate runs asking the participants to either match the positive words with the category “genetic sister” and negative words with the category “adopted sister” or vice versa to match positive words with the category “adopted sister” and negative words with the category “genetic sister”. The order in which the runs are presented counter-balanced across participants and categories switched their localization on the screen in different runs to be on the left or right side of the screen to the same extent. Participants were asked to press a key with either the right or the left hand and thus assign the evaluation word to one category on either the left or right hand side of the computer screen. With the experiment going on, the number of trials in this part of the IAT is increased in order to minimize the effects of practice. The IAT score is based on how long it takes a person to sort the words with the condition associating positive words and genetic (and negative and adopted) in contrast to negative words and genetic (and positive words and adopted). If an implicit preference exist for one of the categories participants would be faster to match positive words to that category relative to the reverse. Data were analysed using Matlab. Similarity between participants’ scores were examined TOST testing ^24^

As a second task, reaction times for the moral decision task were measured with the same group of participants that underwent the IAT. As a difference to the decision task performed during fMRI scanning the order of the decisions was randomized (with easy decision including only strangers and difficult decisions including the sister on one side and the friend on the other). Reaction times were measured as the time between the onset of the slide revealing the identity of the involved individuals and the button press of the participant that related her decision.

### Data availability

The data that support the findings of this study are available on request from the corresponding author MBT. The data are not publicly available due to a prohibition by the Finnish law: Juridical restrictions set by the Finnish law prevent to give public access to the collected data, be it anonymized or non-anonymized data recorded from human individuals. As the consent given by the participants only applies to the specific study reported in our manuscript, no portion of the data collected could be used or released for use by third parties.

## Acknowledgements

Please find here further the Acknowledgement and Contributions of the authors:

Special thanks to Professor Robin Dunbar for the helpful advice on the role kinship in social interactions. We would like to thank Marita Kattelus and Toni Auranen from Advanced Magnetic Imaging (AMI) Centre, Aalto NeuroImaging, Aalto University, Espoo, Finland for their help and support. The data reported in this paper are presented in the Supplementary Methods and archived at the neurovault database: http://neurovault.org/collections/WGSQZWPH/. Financially supported by the Academy of Finland (#276643 and #273469), RFHR (#14-26- 18002), Finnish Cultural Foundation (#150496 to JL)

MBT planning the studies, leading and executing the experimental work, doing the analysis and writing the manuscript

EG and JL developing methods, analysing data and commenting the manuscript

ER assisting in analysing the data and commenting the manuscript

MS and IPJ, planning the studies, supervising the whole project, writing the manuscript

None of the authors claim any conflict of interest. None of the authors reports any competing financial interests.

